# Survival of self-replicating molecules under transient compartmentalization with natural selection

**DOI:** 10.1101/755355

**Authors:** Gabin Laurent, Luca Peliti, David Lacoste

## Abstract

The problem of the emergence and survival of self-replicating molecules in origin-of-life scenarios is plagued by the error catastrophe, which is usually escaped by considering effects of compartmentalization, as in the stochastic corrector model. By addressing the problem in a simple system composed of a self-replicating molecule (a replicase) and a parasite molecule that needs the replicase for copying itself, we show that transient (rather than permanent) compartmentalization is sufficient to the task. We also exhibit a regime in which the concentrations of the two kinds of molecules undergo sustained oscillations. Our model should be relevant not only for origin-of-life scenarios but also for describing directed evolution experiments, which increasingly rely on transient compartmentalization with pooling and natural selection.

## 1. Introduction

Research on the Origins of Life is plagued by several chicken-and-egg problems [1]. One central problem concerns the emergence of functional self-replicating molecules. To be a functional replicator, a molecule must be long enough to carry sufficient information, but if it is too long it cannot be replicated accurately, because shorter non-functional molecules called parasites may replicate faster and take over the system. This was experimentally observed many years ago by Spiegelman [2]. This observation was then rationalized using the notion of error threshold [3], which plays a key role in research on the Origins of Life [4].

Several theoretical solutions have been proposed to address this issue, among which the Stochastic Corrector model [5,6] is prominent. In this model, small groups of replicating molecules grow in compartments, to a fixed final size called the carrying capacity. Then, the compartments are divided and their contents are stochastically partitioned between the two daughter compartments. Thanks to the variability introduced by this stochastic division, and to the selection acting on the compartments, functional replicators can be maintained in the presence of parasites.

Building on this work and on a recent experiment inspired by it [7], we have recently explored alternative scenarios that are also able to maintain information in replicating systems [8,9]. We have proposed a transient compartmentalization dynamics with no cell division, which should be achievable when only prebiotic chemistry is available. In our framework, there are no specific requirements regarding the chemical composition or the topology of the compartment boundaries: transient compartmentalization can result from environment fluctuations due to day-night cycles [10], tides cycles [11] or cycles of confinement and release of chemicals from pores [12].

In this paper, we extend the framework of Ref. [9] to the case of transient compartmentalization of self-replicating molecules. The main new element in this extension is that the selective pressures acting on this system are not externally imposed, as in our previous work, but stem from the system intrinsic dynamics acting as a form of natural selection. Therefore, the assumptions of this new model are agnostic about the environment and its interaction with the system, which is a desirable feature for scenarios on the Origins of Life. Besides, this extension may be also pertinent for certain *in vitro* evolution experiments [13]. Indeed, *in vitro* evolution experiments based on external selection are often more difficult and cumbersome to carry out than the ones based on natural evolution, which we consider here.

In one version of our model, we find oscillatory behavior in the population size of replicators. Although the compartment chemistry we consider has some similarities with hypercycles, which are known to undergo oscillatory behavior in the population size of replicators [14], in our case the oscillations we observe, would not exist in the absence of a transient dynamics of compartmentalization. Thus, our oscillations are more related to the ones reported in experiments using compartmentalized RNA replication systems [15], which also disappear in the absence of compartmentalization. In the Discussion section, we compare the predictions of our model to these experiments and to the theoretical model [16] developed to analyze them. While our framework is applicable to such experimental systems, it is important to appreciate that it has a wide generality. It could equally well describe many other forms of compartmentalized hypercycles or coupled autocatalytic sets, because the self-replicating molecules which we consider need not be RNA replicases.

## 2. Materials and Methods

Here, we introduce two models describing a transient compartmentalization process in which self-replicating molecules (the replicase) may coexist with non-self replicating ones (the parasites) which may be replicated by the first ones. These models are amenable to mathematical analysis. In the first subsection we describe a model of transient compartmentalization where the compartments are populated at each round with an inoculum which has a fixed average size as shown in Figure 1a. In the second subsection, the size of the inoculum is allowed to vary in time as a result of successive dilutions as shown in Figure 1b.

**Figure 1.**
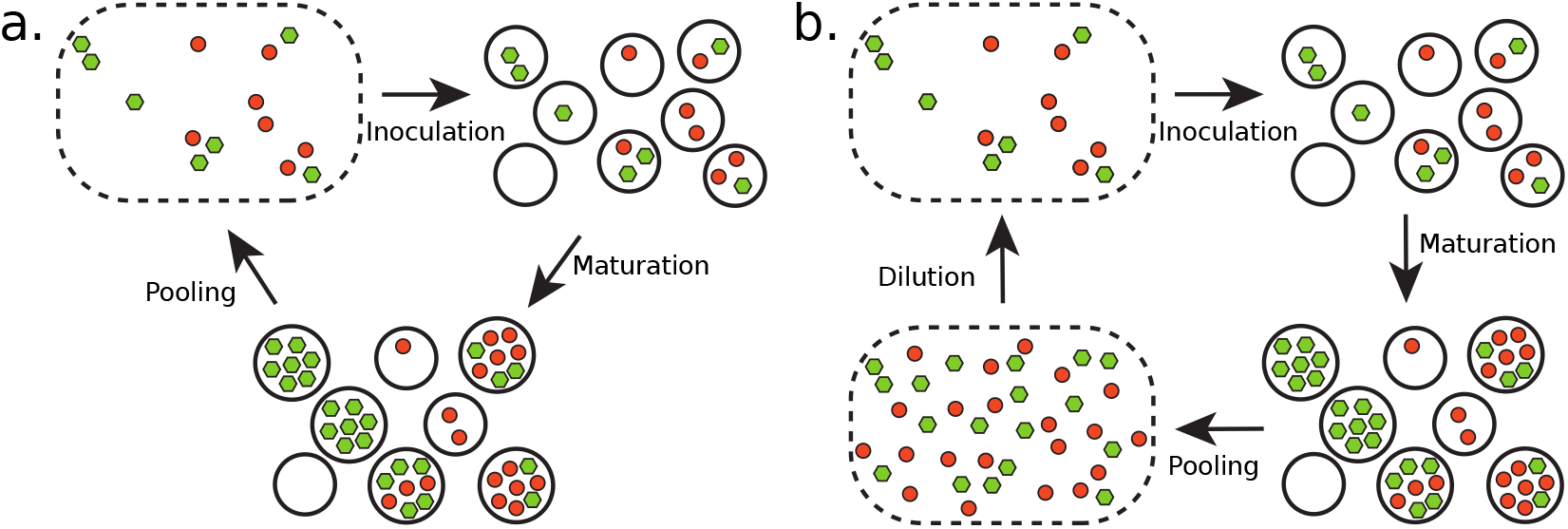
(a) Transient compartmentalization at fixed average number of molecules per compartment, and (b) with a variable average number of molecules. In (a), the cycle splits into steps of inoculation, maturation and then pooling, while in (b) it contains in addition a dilution step. The green and red circles represent the replicators and their parasites respectively.

### 2.1. Transient compartmentalization with a fixed inoculum size

In this subsection, we describe the behavior of a compartmentalized self-replicating system made of two species: self-replicating molecules (replicases) and parasites. Replicases can make copies of themselves and of other parasite molecules, while the parasites can only be copied by the replicases. This is a different case from that discussed in [9], where both the molecules of interests (in that case, the ribozymes) and the parasites could be replicated by externally provided enzymes. In further contrast, in the present case there is no externally applied selection. Thus the main steps of the replicating cycle in this case are as shown in Figure 1a :

- Inoculation of the compartments;
- Maturation of the compartments;
- Pooling of compartment contents.

In the inoculation step, as in Ref. [9], one chooses a number *n* of molecules from the pool, where *n* is Poisson distributed with average *λ*. The resulting inoculum then contains *m* replicases and *y* = *n* − *m* parasites, which are distributed according to a Binomial distribution of parameter *x*, where *x* is the initial fraction of replicases in the pool. We shall denote by *P*_*λ*_ (*n*, *m*, *x*) the resulting probability distribution. This follows closely the corresponding steps in Ref. [9]. However, the dynamics of the maturation step is different and is described by the following equations:

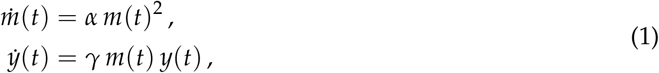

where *m*(*t*) and *y*(*t*) are respectively the self-replicating and the parasitic species populations at time *t*, while *α* and *γ* are their respective replication rates. The analytical solution described in Appendix A yields the compartment composition (*m*(*T*), *y*(*T*)) at the stopping time *T* as a function of the initial composition, denoted by (*m* = *m*(0), *y* = *y*(0)). The stopping time *T* is itself fixed by the condition

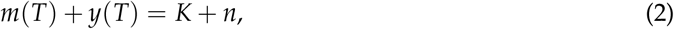

where *n* = *m* + *y* denotes the initial number of molecules in the compartment and *K* is a parameter that represents the number of new strands that can be created during the replication process, due to the finite amount of nutrients present in the compartment. We shall call it the carrying capacity. We shall use in the following the shorthands 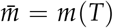 and 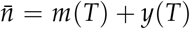. Moreover, the ratio Λ = *γ*/*α* of the replicating constants of both species is another important parameter of the dynamics.

After the maturation step, the contents of the compartments are pooled. The fraction *x*′ of replicases in the pool is expressed in terms of its value *x* at the beginning of the round by the following equation:

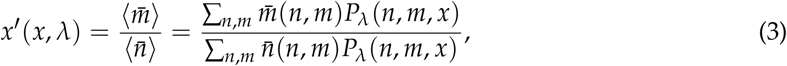

where 〈 · · · 〉 denotes the average with respect to the probability distribution *P*_*λ*_ (*n*, *m*, *x*). Note that the number of molecules in the compartments at the end of the maturation step is not uniform (in particular, compartments which are pure in parasites contain at the end the same number of molecules as in the beginning). Thus we cannot directly average *x* over the compartments as it was done in Ref. [9].

In Appendix B we show that, in the limit Λ ≫ 1 where the parasites are much more aggressive than the self-replicating molecules, the recursion equation (3) can be simplified, yielding

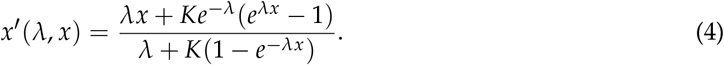

The behavior of this model is described in Section 3.1.

### 2.2. Transient compartmentalization with variable inoculum size

We now consider a model of transient compartmentalization with a variable inoculum size *λ*. In experiments based on serial transfers, a fraction of the solution is transferred into a new fresh medium repeatedly [17]. This can be described theoretically by adding a dilution step in the replicating cycle as shown in Figure 1b. Then *λ* can change because a given amount of the pooling solution can contain a variable number of replicating molecules, depending on their average concentration. The dynamics is now described by a pair of equations for the evolution of the fraction *x* and of the parameter *λ* :

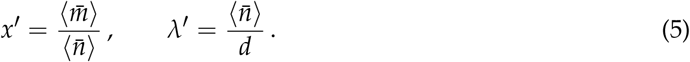

where *d* is the dilution factor and 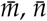 are given by the same equations as above, evaluated with the current value of *λ*.

Using the same approximations used to derive Eq. (4), we obtain the following set of equations, valid for Λ ≫ 1:

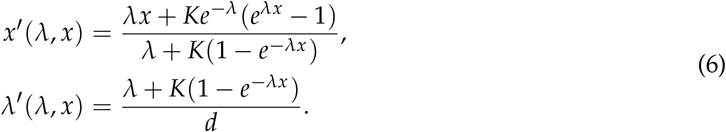

These equations can be more easily manipulated than the corresponding equations for the global compartmentalization process.

The behavior of this model is described in Section 3.2.

## 3. Results

We first describe the results of the model with fixed inoculum size (Figure 1a, Section 2.1), and then those of the variable inoculum-size model (Figure 1b, Section 2.2).

### 3.1. Fixed inoculum size

By studying the stability of the fixed points of the recursion of Eq. (4), we obtain the phase diagram shown in Figure 2, which represent the compositions that are accessible to the system on long times. In contrast to the phase diagram obtained in Ref. [9], we find a large region of coexistence between the self-replicating molecules and the parasites and no pure parasite phase. The absence of the pure parasite phase is expected, since parasites can not grow without replicators. Thus, there are only two phases: a pure replicator phase and a coexistence phase, in which compartments remain of mixed composition. Although the coexistence region appears large, in fact, in the main part of it, self-replicating molecules are maintained at a very small concentration, as shown by the color scale in Figure 2. Therefore, to maintain replicators at a significant concentration, one can not escape the condition that the average size of compartments be of the order of one molecule per compartment.

**Figure 2.**
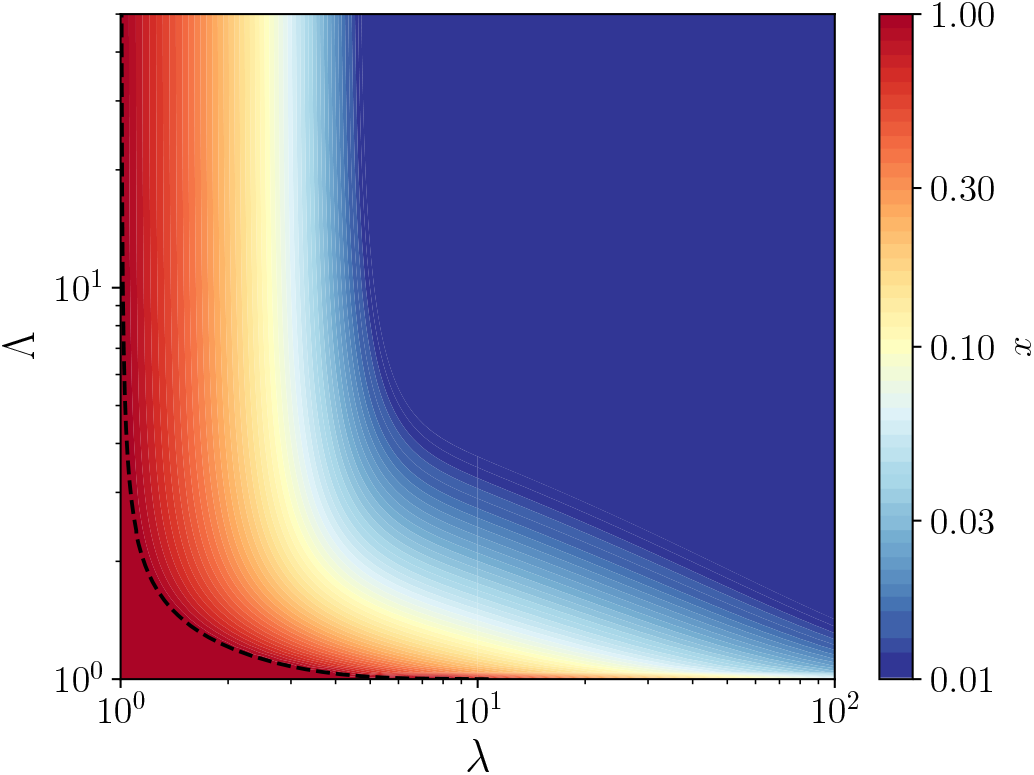
Contour map of the fraction *x* of replicators as a function of (*λ*, Λ), for a carrying capacity *K* = 100, where *λ* denotes the average number of molecules per compartment and Λ the relative growth rates of the parasites with respect to the host. The dotted line is the contour of *x* = 1, which marks the border of the pure replicators phase (the red region). Above this line, a coexistence region exists between the two species at a fraction of replicators indicated by the color scale.

By evaluating the derivative of *x*′ with respect to *x* at *x* = 1, one obtains the equation of the vertical asymptote of the phase diagram separating the pure replicator phase and the coexistence region, which is given by *λ* = 1. The same condition used at *x* = 0 shows that this fixed point is always unstable at finite value of *λ*, which confirms that there is no pure parasite phase. In the coexistence region, a family of vertical asymptotes can be obtained by solving the equation *x*′(*λ*, *x*) = *x* for 0 < *x* < 1 in terms of *λ*. No simple expression has been found for the equations of the corresponding horizontal asymptotes, which separate the pure replicator phase and the coexistence region when *λ* → ∞.

### 3.2. Variable inoculum size

The most striking feature of the model with variable inoculum size, described by equations (5), is the appearance of oscillations in both *x* and *λ*. They are similar to the ones observed in experiments with host-parasite RNAs [15] and modelled numerically in Ref. [16].

When the value of *K* and of *d* are not too large, this system exhibits oscillations in the populations of replicators and parasites shown in Figure 3a. These oscillations can also be seen when representing the fraction *x* of replicators as function of the average compartment size *λ*, as shown in Figure 3b. This behavior can be explained as follows: after the first inoculation, parasites are being replicated quickly by replicators, and the dilution does not counterbalance this increase in population. At some point, the fraction of parasites in the population is so high, that there are not enough self-replicators to contribute to their replication. Then, the dilution has an important effect, since it decreases the population per compartment *λ*, until its average reaches values around *λ* = 1 (see Figure 3b). At this point, compartments contain on average only a single molecule, which can be either a parasite or a self-replicator. The population in empty compartments or compartments containing a single parasite molecule does not grow, therefore only compartments containing a single replicator or containing one replicator and one parasite will contribute substantially to the next round. At this point the replicator population increases, and starts replicating parasites for several rounds, triggering the process again. Note that the mechanism producing these oscillations is different from the Lotka-Volterra one, where the competition between the two species is the main ingredient [18], instead here transient compartmentalization plays an essential role.

**Figure 3.**
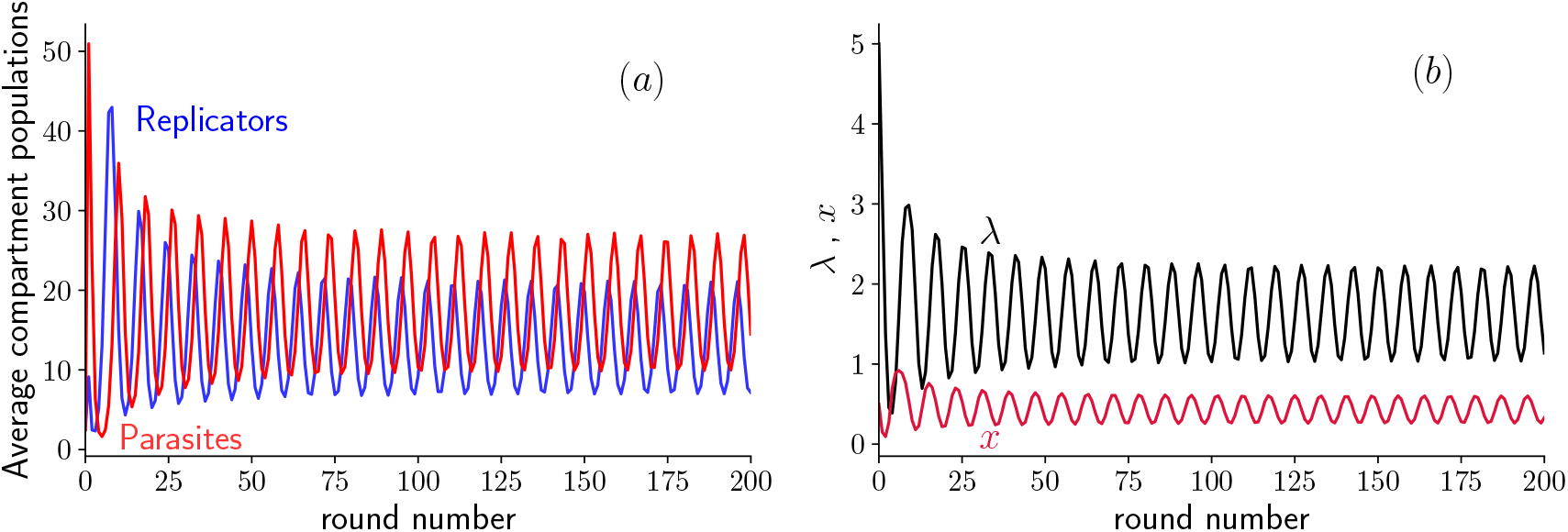
Oscillations in the average amount of self-replicating and parasite molecules per compartment as a function of the round number for *d* = 19, *K* = 60, and Λ = 5. (a): Average population size 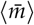 of replicators and 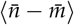 of parasites after the growth step plotted vs. round number. (b): Fraction *x* of replicators and average *λ* of inoculum size. Notice that the oscillations rebound close to the line *λ* = 1.

To delve deeper in the analysis of these oscillations, let us proceed with the equations (6), which are valid in the limit Λ ≫ 1. In a simulation of these equations at a given value of *K*, we observe an abrupt transition when varying the dilution factor. Indeed, when *K* = 60 and *d* = 18, the two average populations oscillate steadily as shown in Figure 4a, while when *d* = 22, oscillations quickly die out as shown in Figure 4b. This abrupt transition is the sign of a bifurcation, which we identify as a supercritical Hopf bifurcation (see Appendix C for more details). The bifurcation occurs at *d* = 20.74 given that *K* = 60. Below this value, the system shows unstable spirals and converges to a limit cycle, while above this value, the system shows stable spirals which converge towards a fixed point (cf. [19, Sec. 8.2]).

**Figure 4.**
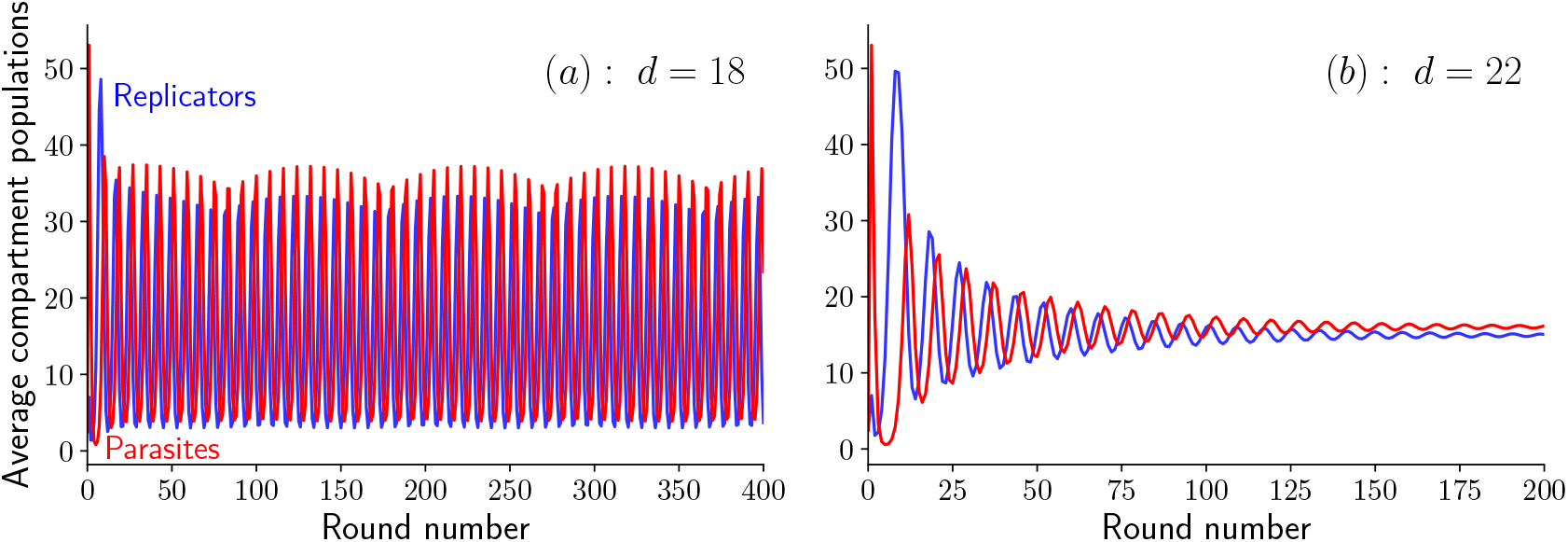
Oscillations in the average amount of self-replicating and parasites molecules per compartment as a function of the round number for *K* = 60 and Λ ≫ 1. (a): Steady oscillations at *d* = 18 (unstable spirals), and damped oscillations at *d* = 22 (stable spirals). Note the beating pattern in the oscillations visible in (a).

When the parameter *d* is further increased still keeping *K* fixed, we find a second transition at *d* = 37.15. At this point, the system no longer oscillates or spirals around a fixed point, but instead converges towards this fixed point monotonically, a case identified as stable node in the literature [19, p. 128].

Another interesting feature in these oscillations is the beating pattern which is visible on Figure 4a as a modulation in the amplitude of the oscillations. This pattern results from the interplay between two frequencies, the sampling frequency fixed by the duration of a single round, and the intrinsic frequency of the oscillations. By changing the sampling frequency, the beating pattern is accordingly modified.

To summarize all these results, we build the phase diagram in the plane (*K*,*d*) shown in Fig. 5a. As can be seen in this figure, the boundaries between the phases, unstable spirals, stable spirals and stable nodes are lines in the plane (*K*,*d*). This can be understood from the following argument. In the limit *K* → ∞, the equations which determine the fixed point coordinates (*x*\*, *λ*\*) deduced from Eqs. 6 can be simplified to yield :

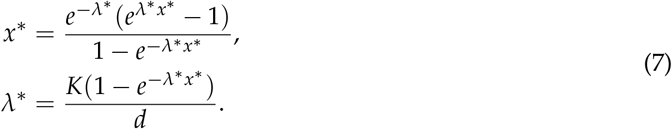

**Figure 5.**
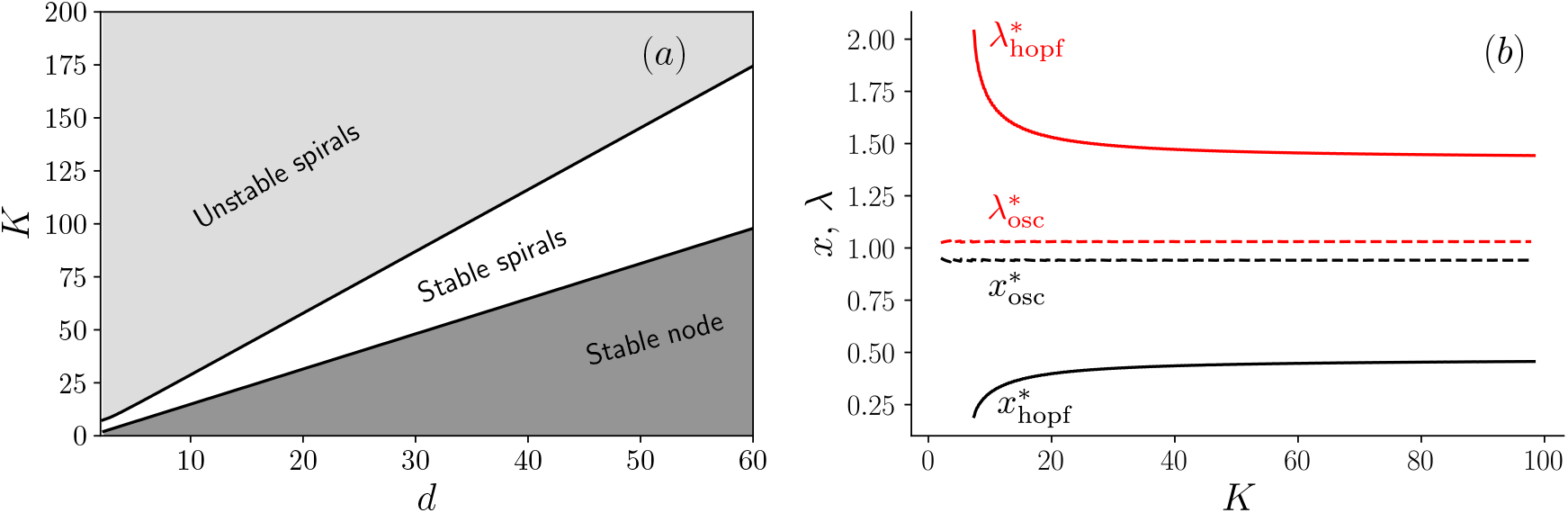
(a) Phase diagram in the plane (*K*,*d*) in the limit Λ ≫ 1 and (b) Evolution of the fixed point coordinates (*x*\*,*λ*\*) as a function of *K*, on the Hopf bifurcation (solid line) and on the transition line between the stable node and stable spirals (dashed line).

The second equation above can be written as

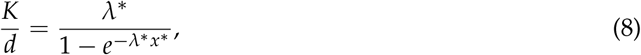

which shows that the coordinates of the fixed point (*x*\*,*λ*\*) only depend on the ratio *K*/*d* in the large *K* limit. It follows that the boundary between the region of unstable and stable spirals, where the Hopf bifurcation occurs is a straight line as shown in Fig. 5a. A similar argument holds for the boundary between the stable spirals and the stable node, which is also a straight line in this diagram.

To confirm this interpretation, we show in Fig. 5b, the values of (*x*\*,*λ*\*) as a function of *K*, evaluated either on the boundary of the Hopf bifurcation and denoted with the subscript ‘hopf’, or on the stable node-stable spirals boundary and denoted with the subscript ‘osc’. When reporting the asymptotic values of (*x*\*,*λ*\*) obtained for large *K* into Eq. (8), one recovers the values of the slopes of two lines in Fig. 5a. In the end, this analysis of the phase diagram in the plane (*K*,*d*) agrees with the general observation made in Ref. [16] that oscillations should be present only in an intermediate range of parameters as far as *K* and *d* are concerned. When either *K* or *d* is very large, one does not expect any oscillations as shown in Fig. 5a.

## 4. Discussion

We have studied a simple system composed of a self-replicating molecule (a replicase) and a parasite molecule that needs the replicase for copying itself. In the case of a fixed inoculum size (i.e., for a fixed value of the parameter *λ*), we have found that this system is able to maintain the replicase molecules against the take-over of parasites in the absence of artificial selection. Although the phase diagram contains a large coexistence region, only in a small part of it, when *λ* is close to one, are the replicase molecules maintained at a significant concentration. This may explain why experiments on directed evolution using compartmentalized self-replicating molecules such as DNA or RNA are usually carried out in this regime for these molecules, while all other required chemical species (nucleotides, other intermediates, …) are typically in excess.

The theoretical framework we have developed here for the case of a variable inoculum size has many similarities with the model proposed in Ref. [16] to explain experiments on host-parasite RNAs [15]. There are however some differences : we consider an infinite number of compartments instead of a finite one, we do not include mutations which could turn the replicase into a parasite, and we do not include local mixing, which means that our model corresponds to the infinite mixing limit of Ref. [16]. Despite these differences, we also find a regime of values of the parameters (in particular for the dilution factor or the carrying capacity) in which sustained oscillations are possible in agreement with Ref. [16]. In Ref. [15], the ratio of catalysis rate constants of the parasite with respect to host, which we denote Λ, was about 5 : this can be obtained by extracting from Table I from that reference, values of *α* = 0.29, *γ* = 1.5 and using Λ = *γ*/*α* ≃ 5 [20]. We find that oscillations are indeed present in our model in this range of the parameters, and oscillation periods comparable to the value reported in Ref. [15] can be recovered from this estimate.

It is interesting to note that both for fixed and variable inoculum sizes, the regime of pure compartments (one molecule per compartment on average) has a particular significance: for a fixed inoculum size, only in this regime can a significant average fraction of self-replicating molecules be maintained, and for a variable inoculum size, only in this regime a rebound can occur in the populations of molecules, allowing oscillations. We surmise that this regime could have a specific significance for the Origins of life. To elaborate a bit on this point, we recall that the emergence of special molecules bearing the genetic information is an essential step in the Origin of Life as emphasized in the RNA world. These molecules are typically found in minority with respect to other species, yet this minority has control of the entire cell [21]. This is a form of information control, which is thought to be one of the key parameters in the Origins of Life [22]. Fluctuations of this minority species therefore have a special role due to their small number. In contrast, many other chemical species, which are not information carriers, are found in large numbers, with fluctuations statistically obeying the law of large numbers. In our model, we see a clear illustration of this mechanism: the replicase behaves as a genome-like molecule, present at the lowest non-zero possible concentration of one molecule per compartment, while all other molecules, which depend on the genome molecules for their own making, are available in the protocell in large concentration.

## 5. Conclusions

Without considering complex chemistry, we have proposed a model which is able to capture important features for Origins of life research, such as the ability to maintain self-replicating molecules using transient compartmentalization and natural selection. An interesting feature of the model with constant inoculum size, is the maintenance of the self-replicating molecule by a form of information control, at the critical level of one molecule per compartment. A striking feature of the model with variable inoculum size, is the appearance of oscillations, which are similar to the ones observed in experiments with compartmentalized self-replicating RNAs [15].

Naturally Ref. [15] has much more than the mere observation of these oscillations. By studying the sequence information of the replicase and its parasites, the authors of that work suggest that parasites can take an active part in the evolution of their host and not just in their own. Different sub-populations of parasites can appear, forming an ecosystem [23] which accelerates evolution. Future studies are needed to quantify these co-evolutionary mechanisms, and perhaps our model could help in that task.

Another important direction for future work would be to consider a large number of interacting chemical species, a situation frequently encountered in Statistical Physics [24]. In this case, we expect that the basic unit of description may no longer be that of single chemical species, but could become collective excitations of the composition, similar to quasi-species [25] or composomes [26]. A general theory of non-equilibrium chemical networks, constrained by conservation laws and symmetries has been recently put forward [27,28]. One attractive feature of such a framework for describing complex chemical systems is that it relies mainly on stoichiometry, therefore the explicit knowledge of the kinetics, which is often missing, is not needed [17].

New types of emergent behaviors could arise by enlarging further the dynamics of compartmentalization. One possibility would be to consider loose compartments [29] or a continuous automated *in vitro* evolution [30]. Besides the relevance for the Origins of Life, we hope that our work could trigger new research directions on applications of transient compartmentalization for chemistry or biochemistry. Perhaps, these new research directions could help overcome practical and fundamental hurdles associated with the synthesis of complex molecules, and facilitate the making of new catalysts or artificial cells [31].

## Author Contributions

Conceptualization, L.P. and D.L.; methodology, G.L.; software, G.L., L. P.; validation, G.L., D.L. and L.P.; formal analysis, G.L.; investigation, G.L.; resources, D.L.; data curation, G.L.; writing–original draft preparation, D.L.; writing–review and editing, L.P. and G.L.; visualization, G.L.; supervision, D.L.; project administration, D.L.; funding acquisition, D.L.

## Acknowledgments

LP is grateful to the Gulliver lab of the ESPCI for a most pleasant hospitality. We would like to thank T. Furubayashi for many insightful discussions and a critical reading of this work. D. L. would like to thank A. Blokhuis for a particularly fruitful collaboration on which this work is built.

## Appendix A. Exact solution of the maturation equations

The maturation equations (1) can be solved analytically to give

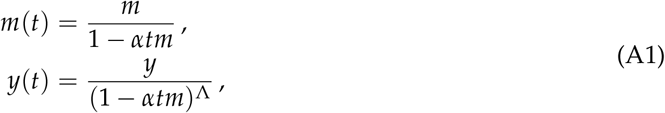

where *m* and *y* are the initial population sizes of both species, *n* their sum, *t* the time, and Λ = *γ*/*α* the ratio of the two replicating constants. The dynamics of these equations is hyper-exponential, and exhibits finite-time divergencies. However, the divergencies are not relevant for the model, since the carrying capacity cutoff stops this dynamics at finite values of *m* and *y*, as intimated by the stopping condition given by Eq. (2). Introducing the quantity *u* = 1 − *αtm*, we express the stopping condition (2) as follows:

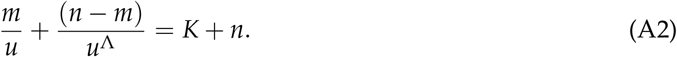

We can then solve this equation in terms of *u* to obtain the final population sizes denoted by 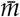 and 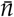.

## Appendix B. Derivation of the equations in the Λ ≫ 1 limit

The expression of *x*′ in the Λ ≫ 1 limit is evaluated by splitting averages in multiple parts. The denominator 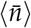 of the recursion (3) is given by

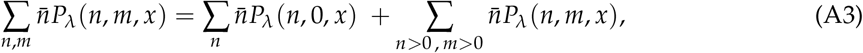

where the first sum of the right hand side corresponds to compartments without self-replicators (*m* = 0) and the second one corresponds to all other compartments containing replicases. Their final populations remain equal to *n* molecules after the maturation in the former compartments without replicators and grow to *K* + *n* molecules in compartment containing replicases. Thus, 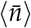 can be expressed by

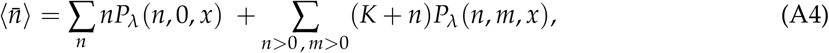

and, using the definition of *P*_*λ*_ (*n*, *m*, *x*) introduced in section 2.1, yields the exact equation

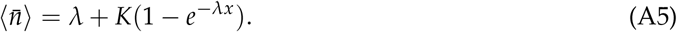

The numerator 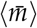 is evaluated in a similar way, starting by splitting the average to give

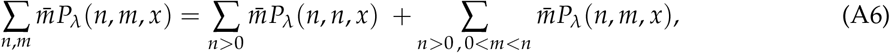

where the first sum of the right hand side corresponds to compartments with only self-replicating molecules (*m* = *n*), and the second sum corresponds to compartments with mixed populations. Empty and pure parasitic compartments do not contribute to the average because in them 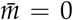. In the former case the final self-replicator population verifies 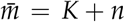. In the latter case, with mixed population, no exact solution for 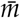 can be determined. We assume that self-replicators do not have the time to replicate during the maturation in presence of aggressive parasites, i.e., that 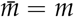 if Λ ≫ 1. Equation (A6) can thus be rewritten

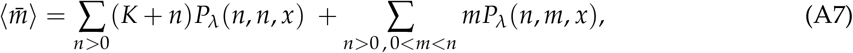

and gives after similar derivations as for equation (A5),

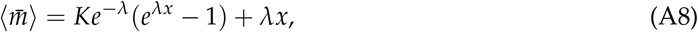

as long as Λ ≫ 1. Finally, by combining equations (A5) and (A8), one obtains the following approximate expression, valid for Λ ≫ 1, of the recursion equation (3):

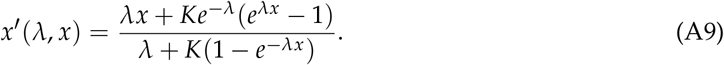

The approximate expression of Eqs. (5) straightforwardly follows.

## Appendix C. Analysis of the bifurcation

In order to analyze the nature of the bifurcation, one could a priori use either the coordinates 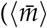, 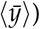 or the coordinates (*x*,*λ*), since there exists a simple bijection between the two sets of coordinates defined by Eq. (5). In the following, we have studied numerically the Jacobian of the system of equations in the coordinates (*x*,*λ*) given by Eqs. (6). We denote the two eigenvalues of this Jacobian by *ϕ_k_* with *k* = 1, 2. The behavior of these quantities is shown in Figure A1. By evaluating the two eigenvalues at the fixed point (*x*\*, *λ*\*), we observe that their modulus moves from above to below 1 as *d* changes from *d* < 20.74 to *d* > 20.74. This means that the fixed point (*x*\*, *λ*\*) with 0 < *x*\* < 1, which is unstable for *d* < 20.74, becomes stable as *d* increases beyond this value. At the transition point for *d* = 20.74, the eigenvalues are complex, which is the indication of a Hopf bifurcation. Below *d* < 20.74, the system spirals up from the unstable fixed point to a stable limit cycle, which encloses the fixed point, while for *d* > 20.74, the system spirals down towards the stable fixed point. We checked that the amplitude of the oscillations decreases smoothly to zero as the bifurcation point is approached, and that therefore the bifurcation is supercritical. There is a second transition at *d* = 37.15, where the imaginary parts of the eigenvalues vanish. This means that the system no longer oscillates or spirals around the fixed point, but instead converges to it monotonically. In this regime, the fixed point is therefore a stable node [19, p. 128].

**Figure A1.**
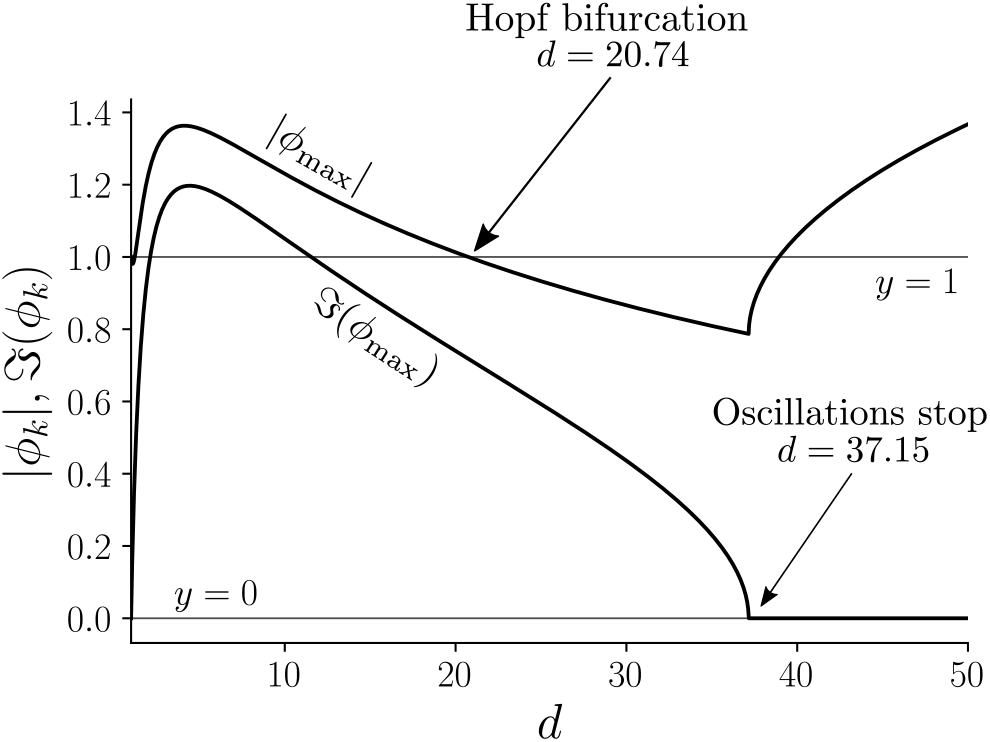
Maximal modulus and maximal imaginary part of eigenvalues of the Jacobian corresponding to Eqs. (6) for a carrying capacity *K* = 60.

